# Physiological concentrations of calcium interact with alginate and extracellular DNA in the matrices of *Pseudomonas aeruginosa* biofilms to impede phagocytosis by neutrophils

**DOI:** 10.1101/2023.10.23.563605

**Authors:** Marilyn J. Wells, Hailey Currie, Vernita D. Gordon

## Abstract

Biofilms are communities of interacting microbes embedded in a matrix of polymer, protein, and other materials. Biofilms develop distinct mechanical characteristics that depend on their predominant matrix components. These matrix components may be produced by microbes themselves or, for infections *in vivo*, incorporated from the host environment. *Pseudomonas aeruginosa* is a human pathogen that forms robust biofilms that extensively tolerate antibiotics and effectively evade clearance by the immune system. Two of the important bacterial-produced polymers in the matrices of *P. aeruginosa* biofilms are alginate and extracellular DNA (eDNA), both of which are anionic and therefore have the potential to interact electrostatically with cations. Many physiological sites of infection contain significant concentrations of the calcium ion (Ca^2+^). In this study we investigate the structural and mechanical impacts of Ca^2+^ supplementation in alginate-dominated biofilms grown *in vitro* and we evaluate the impact of targeted enzyme treatments on clearance by immune cells. We use multiple particle tracking microrheology to evaluate the changes in biofilm viscoelasticity caused by treatment with alginate lyase and/or DNAse I. For biofilms grown without Ca^2+^, we correlate a decrease in relative elasticity with increased phagocytic success. However, we find that growth with Ca^2+^ supplementation disrupts this correlation except in the case where both enzymes are applied. This suggests that the calcium cation may be impacting the microstructure of the biofilm in non-trivial ways. Indeed, confocal laser scanning fluorescence microscopy and scanning electron microscopy reveal unique Ca^2+^-dependent eDNA and alginate microstructures. Our results suggest that the presence of Ca^2+^ drives the formation of structurally and compositionally discrete microdomains within the biofilm through electrostatic interactions with the anionic matrix components eDNA and alginate. Further, we observe that these structures serve a protective function as the dissolution of both components is required to render biofilm bacteria vulnerable to phagocytosis by neutrophils.

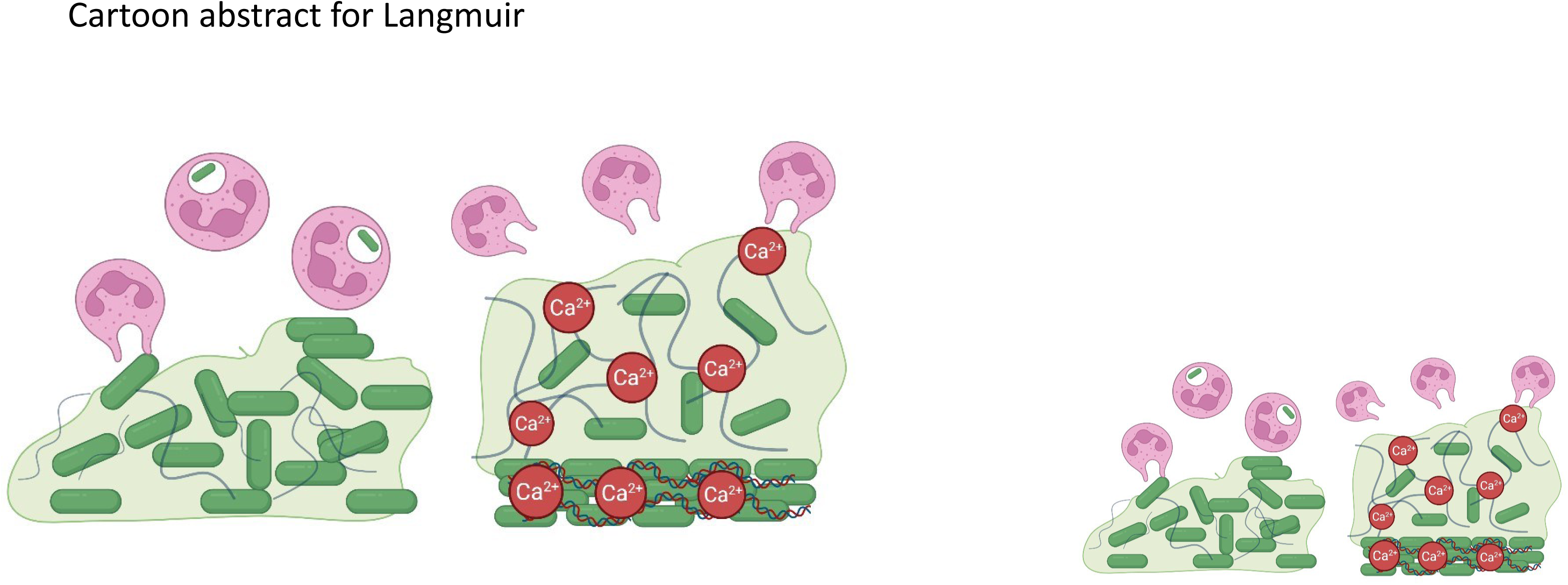

## Introduction

Biofilms are communities of microbes embedded in a matrix of extracellular polymeric substances (EPS), proteins, and other materials. Being in a biofilm protects constituent microbes against treatment with antibiotics and clearance by the immune system. *Pseudomonas aeruginosa* is an opportunistic human pathogen known for producing robust biofilms, notably in chronic wounds and in the airways of persons with cystic fibrosis (CF) [1-3]. An established biofilm infection is difficult to eradicate and is frequently associated with poor patient prognosis in CF [1, 4, 5]. Persistence of chronic *P. aeruginosa* infections in CF patients is largely associated with so-called “mucoid” biofilms that produce high amounts of the matrix polysaccharide alginate [5]. Other matrix components that can be produced by *P. aeruginosa* are the polysaccharides Pel and Psl and extracellular DNA (eDNA). *P. aeruginosa* biofilms have also been shown to incorporate materials from the host environment, such as collagen in the wound bed [6], extracellular DNA from Neutrophil Extracellular Traps [7], and free metal ions including Na^+^, Mg^2+^, and Ca^2+^ [8]. The types and proportions of these matrix components, and the interactions between them, determine the viscoelastic mechanical properties of the biofilm [9, 10].

Recruited via chemical signals to an infection site, neutrophils target pathogens using three primary mechanisms: phagocytosis, degranulation, and NETosis [11]. Neutrophil extracellular traps (NETs) are webs of chromatin that trap pathogens [7, 12-14]. However, bacteria embedded in the matrix are protected from the externally produced NETs. Additionally, the biofilm matrix protects bacteria inside by chemically interacting with substances in the environment, mitigating the diffusion of ions [15] and antibiotics [16-18] and augmenting its structure[19]. Selective diffusion through the matrix not only dampens the penetration of pharmaceutical treatments, but also of neutrophils’ degranulation responses in which antimicrobial, cytotoxic, and proinflammatory materials are released [20]. The combined physicochemical functions of the matrix suggest that mechanical and microstructural disruption of the biofilm are likely important for making the bacteria inside vulnerable to treatment.

Recent work has highlighted the potential importance of biofilm mechanics in impacting clearance in multiple ways, including by the immune system [21-24]. Neutrophils are the most abundant type of granulocyte and play a leading role in the human immune response. Neutrophils effectively phagocytose free-swimming (“planktonic”) bacteria. However, the biofilm structure physically impedes phagocytosis. Aggregates of bacteria in biofilm infections are ∼100 μm in diameter, whereas neutrophils are ∼10 μm in diameter. When challenged with engulfing an unyielding, rigid target larger than its diameter, a neutrophil exhibits “frustrated phagocytosis”, in which the cell spreads itself on the target but fails to close and form a phagosome, potentially releasing toxic and inflammatory materials which further harm the host [14]. New treatment approaches have focused on altering biofilm viscoelasticity [19, 21, 22, 25] by understanding the role of individual matrix components. Our recent studies of hydrogel models for biofilm mechanics have shown that bulk rheological properties, namely elastic modulus and toughness, are negatively correlated with phagocytic success by neutrophils [26, 27]. Additionally, we have shown that treatment with polymer-degrading enzymes can significantly alter the mechanics of biofilms grown *in vitro* [28]. Therefore, we expect that mechanical compromise by enzymatic treatment should promote phagocytic success. However, biofilms have a highly heterogeneous internal microstructure. Bulk mechanics alone is insufficient to describe the importance of local micro-scale structure and the contribution of each EPS component to the recalcitrance of biofilms.

Here we investigate the microstructure and mechanics of mucoid biofilms, grown without and with calcium, and the mechanical impact of targeted enzyme treatment thereon; phagocytic clearance of biofilm bacteria by neutrophils is measured under all growth conditions and enzyme treatments studied. We grow alginate-overproducing biofilms *in vitro* from *P. aeruginosa* lab strains in the presence of physiologically-relevant Ca^2+^ concentrations and treat them with targeted enzymes alginate lyase and DNAse I. To quantify the mechanical response of these biofilms to enzymatic treatment, we measure changes in relative viscoelasticity using passive multi-particle tracking microrheology. We use scanning electron microscopy (SEM) and confocal laser scanning microscopy (CLSM) to qualitatively assess the structural features of treated and untreated calcium-gelled biofilms and observed the prominent presence of eDNA in a distinct regional architecture. We further investigate the ability of neutrophils to phagocytose bacteria from these biofilms using a pH-dependent stain and CLSM.

Our findings suggest that the role of eDNA in biofilm architecture is augmented by the presence of divalent cations. For biofilms grown with Ca^2+^, combined treatment with both alginate lyase and DNAse I is necessary to increase the vulnerability of biofilm bacteria to phagocytosis by neutrophils. This work shows the need to consider both micro-architecture and viscoelasticity when developing biofilm treatment methods.

## Materials and Methods

### Isolation of neutrophils

Neutrophils from adult volunteer blood donors were isolated according to published methods [27]. Whole blood was collected into 10mL lithium-heparin coated tubes (BD Vacutainer, Franklin Lakes, NJ) by venous puncture. Blood was mixed at a 1:1 ratio with a filter-sterilized solution containing 3% dextran and 0.9% sodium (Sigma-Aldrich, Burlington, MA). Red blood cells sedimented and fell to the bottom of the tube. The supernatant was removed and centrifuged (Eppendorf 5810R, A-4-62 Rotor, Enfield, CT) for 10 minutes at 500 x g, separating blood cells from plasma. The resulting pellet of cells was resuspended in Hanks Buffered Salt Solution (HBSS) without calcium or magnesium (Gibco Laboratories, Waltham, MA) and gently placed onto 4mL of Ficoll-Paque density gradient (GE Healthcare, Chicago, IL). This gradient was centrifuged for 40 minutes at 400 x g. The supernatant was discarded and remaining red blood cells were lysed by suspending the pellet in 4mL deionized water for 30 seconds. 4mL of filter-sterilized 1.8% NaCl was added to restabilize the solution. After 5 minutes centrifugation at 500 x g, the pellet of neutrophils was buffered in 800 μL HBSS with calcium and magnesium (Gibco Laboratories, Waltham, MA) and 200 μL human serum (Sigma, St. Louis, MO). One 10 mL tube of whole blood yielded 1 mL of neutrophil-only solution. If more than one tube of blood was drawn, the resulting suspensions were mixed thoroughly by pipette prior to experimental use.

### Growth of mucoid biofilms

The bacterial strain used in this study was a mucoid (alginate-overproducing) laboratory strain of *P. aeruginosa*, PA01 Δ*mucA.* Glycerol bacterial stock stored at -80 °C was streaked onto sterile Luria-Bertani (LB) (Fisher, Waltham, MA) agar plates and grown overnight at 37 °C. One colony was selected, cultured in 4mL of LB broth liquid media (Fisher, Waltham, MA), and grown shaking overnight at 37 °C. Biofilms without Ca^2+^ were grown by spreading 250 μL of overnight culture onto LB agar plates, and incubating statically overnight. To grow biofilms with 5mM Ca^2+^, 5% CaCO3 (Sigma, St. Louis, MO) and 5% glucono-δ-lactone (GDL, Sigma, St. Louis, MO) were combined with the overnight culture before spreading onto agar plates.

### Quantification of bacterial cell count in biofilms

Biofilms were grown in 96-well plates (Corning, Corning, NY) by adding 250 μL of raw culture or culture supplemented with 5mM Ca^2+^ according to the recipe in the above section. Biofilms were allowed to grow overnight statically at 37 °C. A serial dilution was performed, and 60 μL of diluted solution was spread onto fresh LB agar plates with an L-spreader. These were allowed to grow overnight, and colony-forming units (CFUs) were manually counted the following day. Experiments were performed in biological triplicate and results shown in Table S1. No statistically-significant difference in the cell ounts of biofilms grown without and with calcium was found.

### Targeted enzyme treatment

Solutions of alginate lyase were prepared by dissolving the enzyme in its powdered form (Sigma, St. Louis, MO) into deionized water at a concentration of 200 U/mL. Solutions of Optizyme DNAse I (ThermoFisher, Waltham, MA) were prepared with DNAse I Buffer Solution (ThermoFisher, Waltham, MA) at a concentration of 500 U/mL. DNAse I is buffered in 100mM Tris-HCl, 25mM MgCl2, and 1 mM CaCl2. The above enzyme concentrations are used throughout the study. Biofilmswere gently scraped from agar plates and placed in either glass-bottom petri dishes (Matsunami, Bellingham, WA) for microrheology experiments or in wells of a 24-well plates for phagocytosis assays. As such, samples of the same biofilm were used for both measurements. For microrheology measurements, they were then treated with 2 μL of enzyme, submerged in fresh LB broth to prevent dehydration, and incubated statically at 37° C for 1 hour. For phagocytosis assays, biofilms were treated with 12.5 µL of enzyme and incubated for 1 hour. For both microrheology and phagocytosis measurements, the control experiment was done by adding the corresponding volume of the appropriate buffer without the enzyme to the biofilm sample.

### Scanning electron microscopy analysis of biofilm structure

Biofilms were grown on squares of ACLAR® (Ted Pella, Redding, CA) substrate, placed in 24-well plates (Corning, Corning, NY), and incubated with 200 µL neutrophil solution (See *Isolation of neutrophils*) for 1 hour at 37° C. Samples were then fixed for microscopy following enhanced biofilm fixation methods [29]. In brief, biofilms were fixed overnight with 4% Glutaraldehyde (Electron Microscopy Sciences, Hatfield, PA), 2% paraformaldehyde (Electron Microscopy Sciences, Hatfield, PA), and 0.15% alcian blue (Sigma, St. Louis, MO) in 0.1M Na Cacodylate buffer of pH 7.4 (Electron Microscopy Sciences, Hatfield, PA). Samples were washed thrice with the same buffer solution, and post-fixed in a solution of 1% osmium tetroxide (Ted Pella, Redding, CA) and 1% tannic acid (Electron Microscopy Sciences, Hatfield, PA). After 2 hours, the samples were dehydrated using graded ethanols (ThermoFisher, Waltham, MA) and hexamethyldisilazane (HMDS, Ted Pella, Redding, CA), then sputter-coated with 12 nm platinum/palladium with a Cressington 208HR sputter coater (Ted Pella, Redding, CA). Scanning electron microscopy (SEM) was performed under high vacuum with a Zeiss Supra field emission SEM using an SE2 detector and a 5kV accelerating voltage. False-coloring of images was performed with GNU Image Manipulation Program (GIMP).

### Assessment of eDNA presence using confocal laser scanning microscopy

Biofilms were grown in glass-bottom dishes (Matsunami, Bellingham, WA) by adding 250µL of culture with appropriate concentrations of CaCO3 and GDL (see *Growth of mucoid biofilms*) and allowed to incubate statically overnight at 37°C. Biofilms were washed thrice with fresh phosphate buffered saline (PBS, Sigma, St. Louis, MO) before being stained with 200 µL of Propidium Iodide (Calbiochem, San Diego, CA) buffered in PBS at a concentration of 1mg/mL. Samples were incubated for 20 minutes at 37°C and subsequently visualized with confocal laser scanning microscopy (Olympus IX71 inverted confocal microscope, 60X oil-immersion objective) using FluoView FV10-AWS version 04.02 from Olympus America. Z-stacks were recorded and 3D renderings reconstructed with ImageJ (Fiji) [30].

### Passive multi-particle tracking microrheology

For multi-particle tracking, videos of each sample were captured using an Olympus IX71 phase contrast microscope and OrcaFlash4.0 v2 camera, at a frame capture rate of 33 fps. Tracer beads (Bangs Laboratories, Fishers, IN) embedded in samples were tracked over the course of an 8 second video. The time-dependent displacements of tracer particles can be analyzed using the Generalized Stokes-Einstein Relation to determine rheological properties of the material [31, 32]. Passive microrheology depends on the thermal fluctuations (on the order of kBT) to displace the particles along random trajectories. The particles’ mean square displacement (MSD) represents strain of the material caused by thermal stress. The relative viscoelasticity, 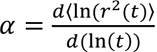, is given by finding the slope MSD versus lag time on a log-log plot. This value ranges from 0 to 1, corresponding to an elastic solid and viscous fluid, respectively [33]. Intermediate values indicate how relatively elastic or relatively viscous the material is. All microrheology experiments were each done on three biological replicates, with 4-5 technical replicates per biological replicate. The number of tracks reported for each sample varies and is listed in Table 1.

**Table 1.** Number of particle tracks analyzed for each treatment condition.

|  | Control | Alginate Lyase | DNase I | Both |
| --- | --- | --- | --- | --- |
| 0mM $\text{Ca}^{2+}$ | 88 | 77 | 87 | 65 |
| 5mM $\text{Ca}^{2+}$ | 92 | 88 | 123 | 59 |

### Data analysis

Tracer particles were tracked across the sequence of images using the Fiji [30] installation of ImageJ and the TrackMate plugin [34]. Centroids of particles are located using a Laplacian of Gaussian filter, and 2D tracks are generated by linking particle positions from frame to frame with a Simple Linear Assignment algorithm. The resultant tracks are input to MATLAB and analyzed using msdanalyzer [35], a MATLAB class for tracking particle trajectories and their resultant mean square displacements. Student T-tests and single-factor ANOVA tests for statistical significance were performed in Microsoft Excel.

### pH-dependent stain phagocytosis assay

Treated biofilms were submerged in 500 μL PBS and stained with 0.5 μL of pHrodo Red, succinimidyl ester (ThermoFisher, Waltham, MA) from a 1mg/150μL stock solution in DMSO (Sigma, St. Louis, MO) and incubated at room temperature for 30 minutes. The dye was removed and biofilms washed thrice with PBS. 200 μL of neutrophil suspension (see *Isolation of neutrophils*) was added to each well and allowed to incubate at 37°C for 30 minutes. Neutrophils were then removed and analyzed using confocal laser scanning microscopy (Olympus IX71 inverted confocal microscope, 60X oil-immersion objective) using FluoView FV10-AWS version 04.02 from Olympus America. At least 100 neutrophils per sample were counted and assessed for phagocytic success as indicated by red fluorescence within the phagosome. Student T-tests and single-factor ANOVA tests for statistical significance were performed in Microsoft Excel. The total number of neutrophils counted for each treatment type and the corresponding significances are shown in Table 2.

**Table 2.** Statistics for phagocytosis assays.

| **0mM Ca<sup>2+</sup>** |  | Control | Alginate Lyase | DNase I | Both |
| --- | --- | --- | --- | --- | --- |
| Biological Replicates | N=4 |  |  |  |  |
| ANOVA p-value | 0.00161 |  |  |  |  |
| Total neutrophils counted |  | 446 | 528 | 486 | 492 |
| Student t-test p-value (relative to control) |  |  | .00989 | .29886 | .00655 |
| **5mM Ca<sup>2+</sup>** |  |  |  |  |  |
| Biological Replicates | N=3 |  |  |  |  |
| ANOVA p-value | 0.00366 |  |  |  |  |
| Total neutrophils counted |  | 462 | 398 | 336 | 332 |
| Student t-test p-value (relative to control) |  |  | .46235 | .20066 | .00302 |

Ethical statement: Work with neutrophils was approved by the Institutional Review Board at the University of Texas at Austin (Austin, TX) as Protocol No. 2021-00170.

## Results and Discussion

### For mucoid biofilms grown without Ca^2+^, treatment with alginate lyase increases the phagocytic success of neutrophils

Alginate consists of β-D-mannuronate and α-L-guluronate monomers, which can be homogenously or heterogeneously linked into longer chains [36]. Alginate lyase degrades alginate via β- elimination reaction, which breaks down glycosidic bonds between monomers. Alginate lyases have been used in treatment of CF to enhance the efficacy of antibiotics [37-39].

In the absence of added Ca^2+^, when mucoid biofilms grown from the lab strain PA01 Δ*mucA* are treated with alginate lyase, we find that 62% more neutrophils successfully engulf bacteria than in the control case, in which the biofilm is not treated by enzyme (Figure 1A). This result is consistent with alginate providing a protective barrier that can be compromised by degrading polymers into smaller units. To determine the relative contribution of eDNA to protection from neutrophils, we also treat biofilms with DNAse I and with a combination of DNAse I and alginate lyase. DNAse I alone has no statistically-significant effect, suggesting that in these biofilms eDNA does not play a substantial role in inhibiting phagocytosis (Figure 1A). The combination treatment yields similar results to that of alginate lyase alone (Figure 1A). The hydrodynamic radius of DNAse I, based on its molecular weight of ∼30 kDa, can be approximated to be in the range of 2-3 nm [40]. The mesh size of the *P. aeruginosa* biofilm matrix has been approximated to be on the order of 100 nm [41, 42]. Therefore, it is likely, although not certain, that the inefficacy of DNAse I is not due to inability to diffuse through the matrix. These results indicate that alginate is the dominant EPS polymer hindering phagocytic engulfment when growth media is not supplemented with Ca^2+^.

**Figure 1.**
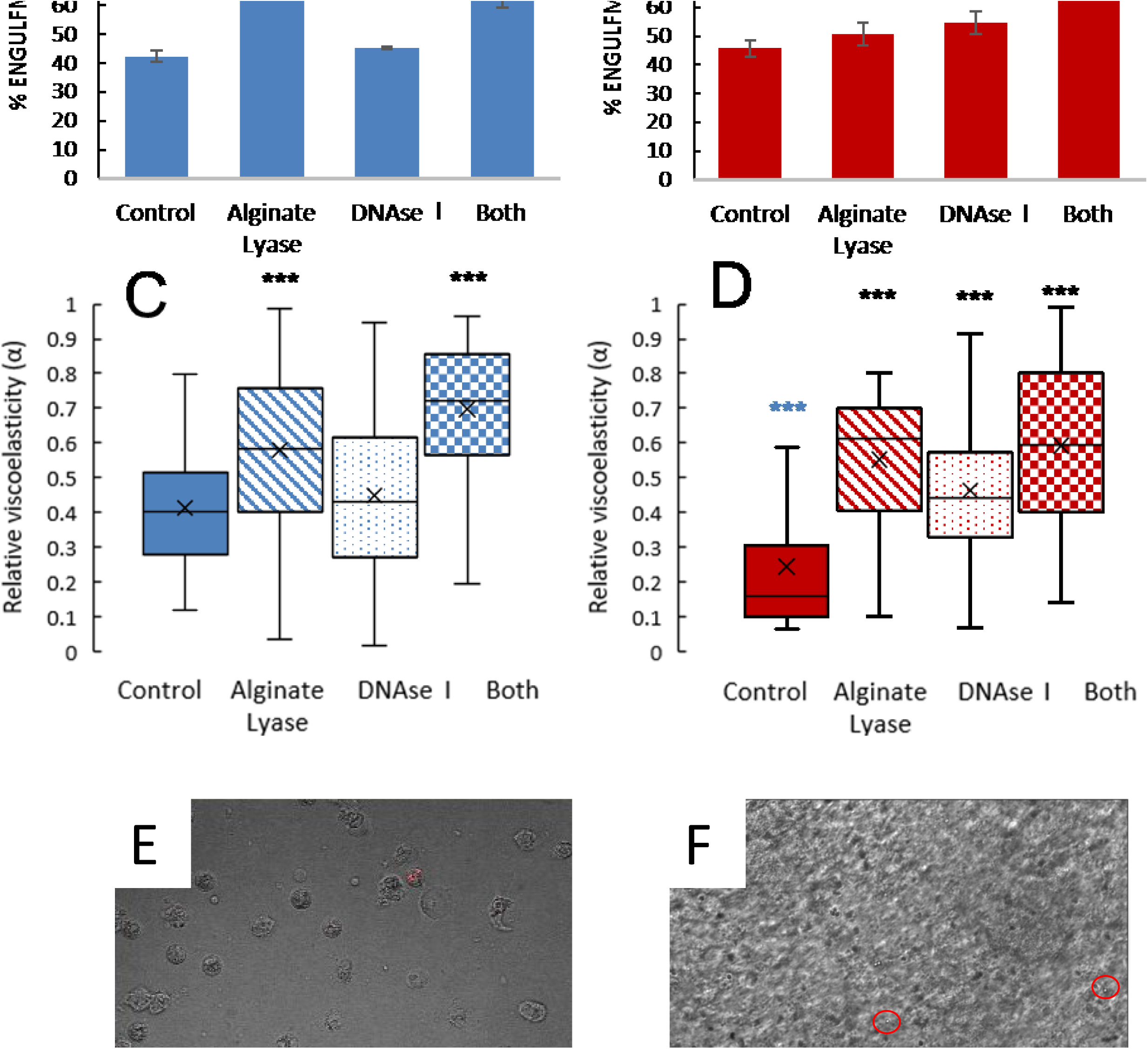
(A,B) The effects of enzyme treatments on neutrophil engulfment success rates. (C,D) relative viscoelasticity measurements for mucoid biofilms grown without (blue) and with (red) 5mM Ca2+. (E) Representative image of neutrophils after phagocytosis assay. Red fluorescence of pHrodo indicates successful phagocytosis. (F) Representative still image of micro-beads (circled in red) embedded in a biofilm during microrheology experiments. Box-and-whisker plots represent data taken across all biological replicates. Boxes show the middle 50% of the data, with horizontal lines and x-markers representing median and mean, respectively. Upper/lower whiskers represent upper/lower quartiles. Black statistical markers (all panels) indicate a statistically-significant difference of enzyme-treated samples from their respective controls. Blue statistical markers (panel D) indicate a significant difference between controls of the two groups. ** p<.01, *** p<.001. Scale bars represent 20 μm

### For mucoid biofilms grown with Ca^2+^, treatment with a combination of both alginate lyase and DNAse I is required to increase the phagocytic success of neutrophils

Ca^2+^ is present in the body as a free ion concentration in the range of 1-5mM. Such concentrations of calcium ions have been shown to form ionic bonds with alginate, thus producing a hydrogel-like biofilm structure with a diffuse arrangement of bacterial cells [8, 43, 44]. Ca^2+^ also plays an important role in the structural integrity of a biofilm by electrostatically binding with eDNA to form bacterial aggregates [45]. Thus, calcium-induced changes to EPS due to interactions with alginate and eDNA have been explored individually [12, 44-46], but the simultaneous bulk and micro-scale effects when both alginate and eDNA are prominent have not been well-characterized.

For biofilms grown with a physiological concentration of 5mM Ca^2+^, treatment with alginate lyase does not significantly increase phagocytic success (Figure 1B). DNAse I also has little effect independently. A combination of both enzymes is the only tested treatment which promotes significantly greater engulfment success, 68% greater than the control (Figure 1B). Thus, we find that growing biofilms with a physiological concentration of 5mM Ca^2+^ radically changed the effect of enzymatic treatment on phagocytic success, compared with biofilms grown without Ca^2+^.

### Treatment with alginate lyase increases the relative viscosity of mucoid biofilms (grown without Ca^2+^)

To assess the changes in biofilm mechanics caused by enzymatic treatment, we perform multi-particle tracking passive microrheology to measure relative viscoelasticity (α). Values of 0 and 1 correspond to a purely elastic solid and a purely viscous fluid, respectively. Ensemble averaged MSD-lag time plots for each biofilm type and treatment combination are shown in Supplementary Fig. S1. We observe a significant increase in α for mucoid biofilms treated with alginate lyase, indicating a shift toward being more relatively viscous (Figure 1C). This suggests that alginate structures have been effectively degraded to render the biofilm more fluid-like. The effect of DNAse I on α is insignificant, and the combination treatment yields similar changes to that of alginate lyase alone (Figure 1C). Taken together with the data in Figure 1A, our results indicate that a shift toward relative viscosity is associated with an increase in phagocytic success for these biofilms grown without added Ca^2+^. Our results also indicate that, under these conditions, alginate is likely the primary EPS component associated with solid-like (elastic) properties and mechanics-based immune evasion of mucoid biofilms; eDNA does not appear to contribute in a substantial way.

### Physiologically-relevant Ca^2+^ concentration increases relative elasticity and the role of eDNA

To analyze the changes in biofilm mechanics caused by growth with Ca^2+^ and the subsequent changes in biofilm mechanics following enzymatic treatment, we use passive particle-tracking microrheology to measure relative viscoelasticity (α). We find that biofilms grown with Ca^2+^ are significantly more relatively elastic than are biofilms grown without calcium (Figure 1C,D). All three tested enzyme treatments resulted in a significant shift toward relative viscosity (Figure 1D). This finding indicates that both alginate and eDNA may play a substantial role in biofilm structure when calcium ions are present. Like alginate, eDNA carries a negative charge, so ionic bonding with cationic Ca^2+^ is expected [45]. However, the lack of correlation between phagocytic success and relative viscoelasticity (Figure 1 B,D) suggests a more complex immune evasion mechanism than mechanics *per se*.

### *Growth with* Ca^2+^ *increases the amount and spatial heterogeneity of eDNA within the biofilm matrix*

To assess the distribution of eDNA within the biofilm matrix, we stained biofilms (grown from GFP-expressing bacteria) with propidium iodide and imaged them using confocal laser scanning microscopy. Three-dimensional reconstructions reveal that, in biofilms grown without Ca^2+^, DNA (shown in red in Figure 2) is present in small, discrete regions when no calcium was added (Figure 2A,C). In contrast, when the biofilm has been grown with 5mM Ca^2+^, biofilms are thicker and more eDNA is present (Figure 2B). Moreover, the DNA forms a dense, contiguous layer (Figure 2D), likely due to crosslinking by Ca^2+^.

**Figure 2.**
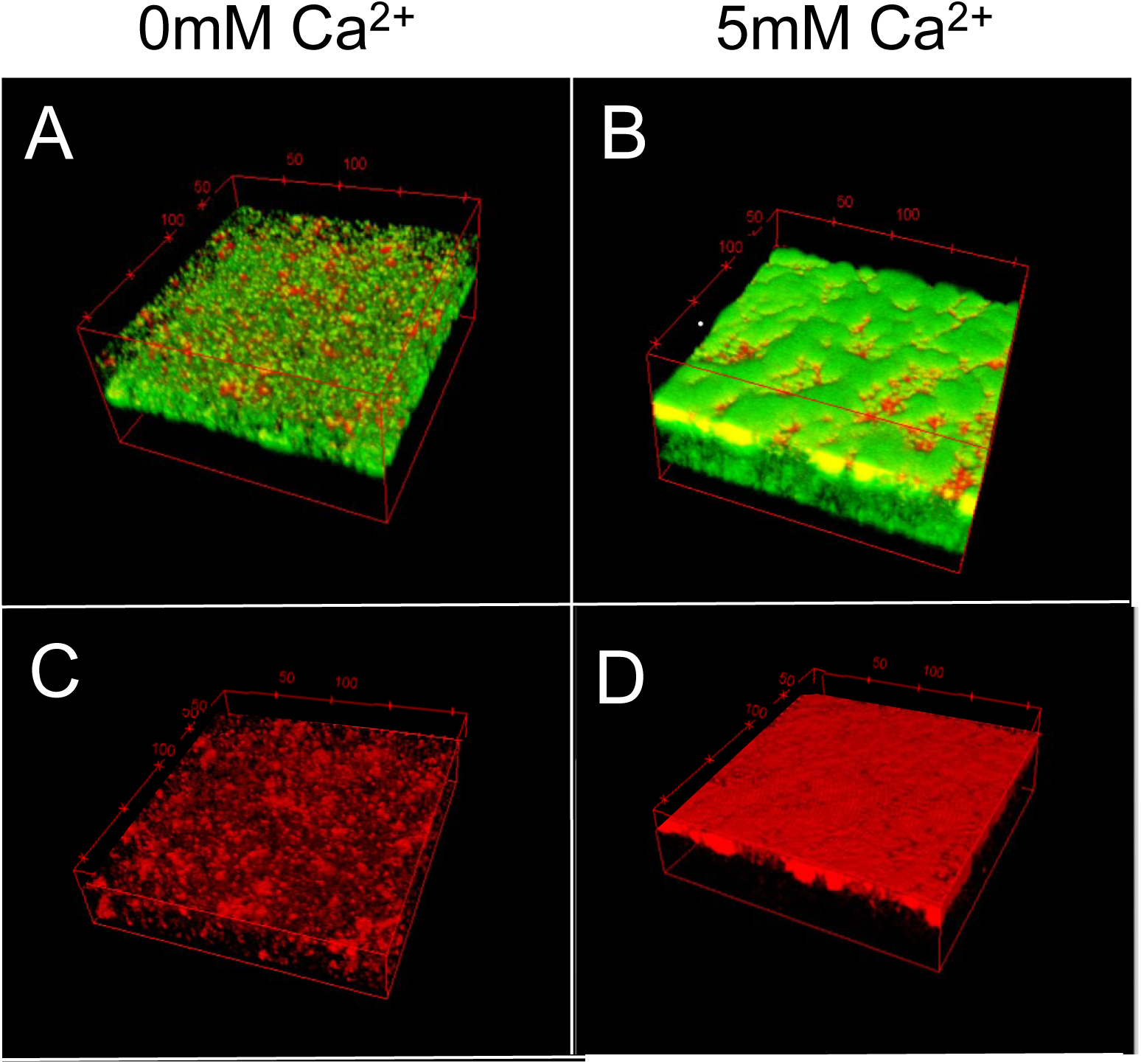
Confocal laser scanning microscopy 3D reconstructions of biofilms grown without Ca ^2+^ (A) and with 5mM Ca^2+^ (B). GFP fluorescence of biofilm biomass is shown in green, and DNA stained with propidium iodide is shown in red. Panels C and D show renderings of DNA fluorescence only. Renderings of biomass fluorescence only are provided in Supplementary Fig. 2. Coordinate boxes have dimensions of 212×212×100µm.

Our results are consistent with previous reports [44] in which increased thickness is attributed to a more diffuse distribution of bacterial cells within the gelled biofilm. The cause of enhanced DNA presence, however, is not clear. We speculate that one or more of the following might apply. Ca^2+^ may induce the release of eDNA, though reports of this effect suggest a species-and strain-dependence [6, 31, 32]. Ca^2+^ has also been shown to enhance adhesion of bacterial cells to alginate films [47], as well as enhance the adsorption of eDNA to surfaces and increase the work required to remove eDNA [48]. Thus, the increase in eDNA presence may also be a consequence of more efficient surface colonization and biofilm formation. Established and strongly attached biofilms are also more likely to withstand the washing procedure associated with staining and imaging, which may contribute to the observed increase in thickness.

For biofilms grown with 5mM Ca^2+^, we propose that the inefficacy of either alginate lyase or DNAse I alone at increasing engulfment success rates (Figure 1B) may be a consequence of the dense, extensive layer of eDNA shown in Figure 2 B and D.

### Scanning electron microscopy reveals shield-like heterogeneities for biofilms grown with Ca^2+^

To investigate this Ca^2+^-dependent, eDNA-based planar layer at higher resolution, we preserve biofilms using fixation methods detailed in [29] and image the fixed biofilms with scanning electron microscopy. Figure 3 shows representative images of untreated (top row) biofilms grown without (3A) and with (3B) added Ca^2+^. Panels C and D show the corresponding biofilm treated with alginate lyase. A treated biofilm grown without added Ca^2+^ is shown in Figure 3C. In striking contrast, in biofilms grown with Ca^2+^, we observe a sheet-like architecture of closely-packed, aligned bacterial cells underneath a thick layer of EPS that we presume is predominantly alginate (Figure 3D). These images agree with the layered architecture observed in Figure 2, where a dense layer of eDNA was verified by propidium iodide staining.

**Figure 3.**
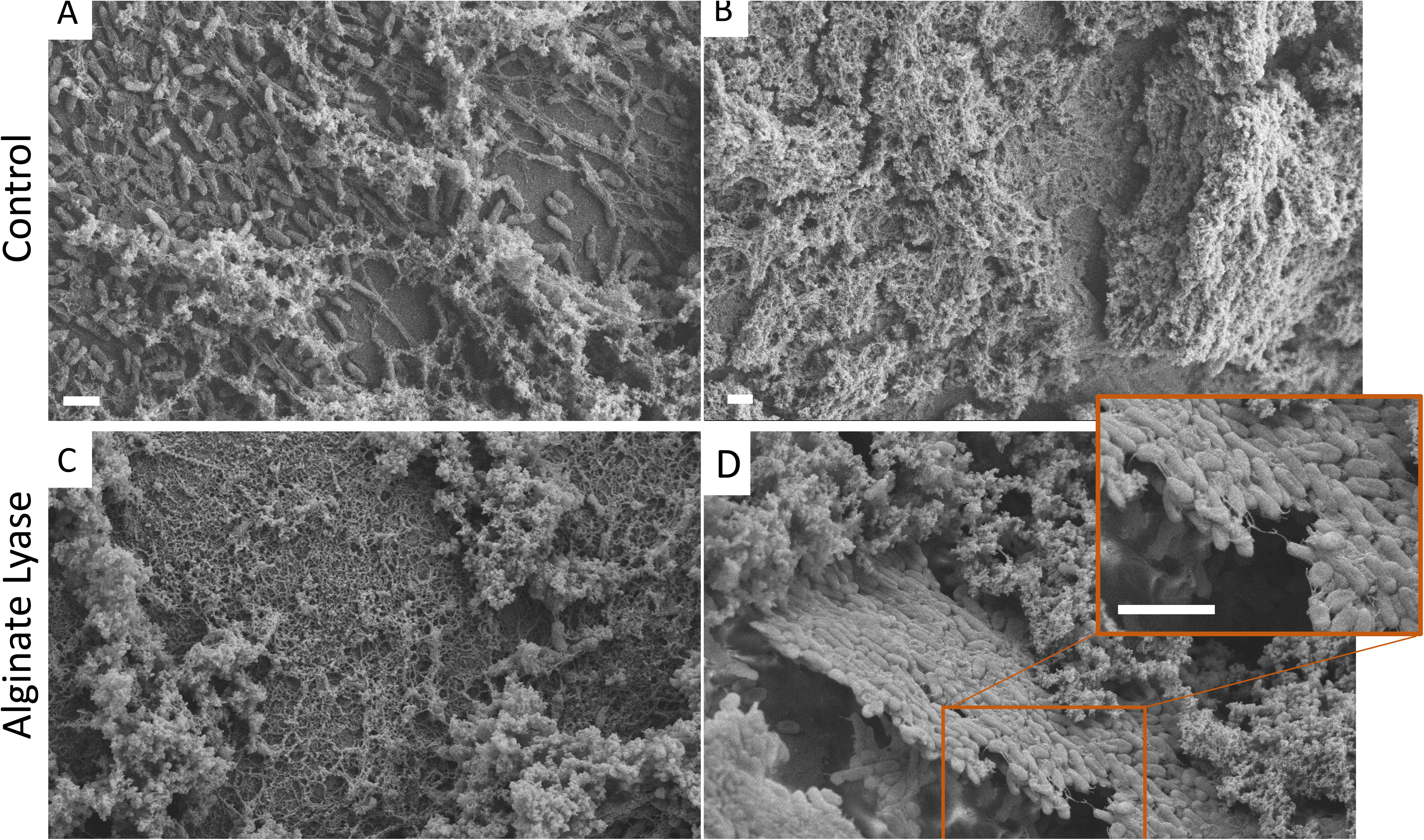
Representative SEM images of untreated mucoid biofilms grown without (A) and with (B) the addition of 5mM Ca^2+^. Panels C and D show corresponding biofilms which have been treated with 200 U/mL alginate lyase, revealing the underlying structure. Scale bars represent 2 µm. The fluffy material shown in all panels is the EPS component alginate. The inset in Panel D shows a higher magnification of the ‘shield-like’ structure of bacterial cells bound together by fibers of eDNA. For ease of distinguishing between features, the image in Panel D has been reproduced in false coloring as Supplementary Fig. 3.

To gain further insight into neutrophil interactions with enzyme-treated biofilms treated and the impact of biofilm microstructure thereon, we image samples corresponding to the treatment conditions in Figure 1A and 1B. All biofilms have been incubated with freshly isolated neutrophils for 60 minutes prior to fixation. Representative images of our results are shown in Figure 4.

**Figure 4.**
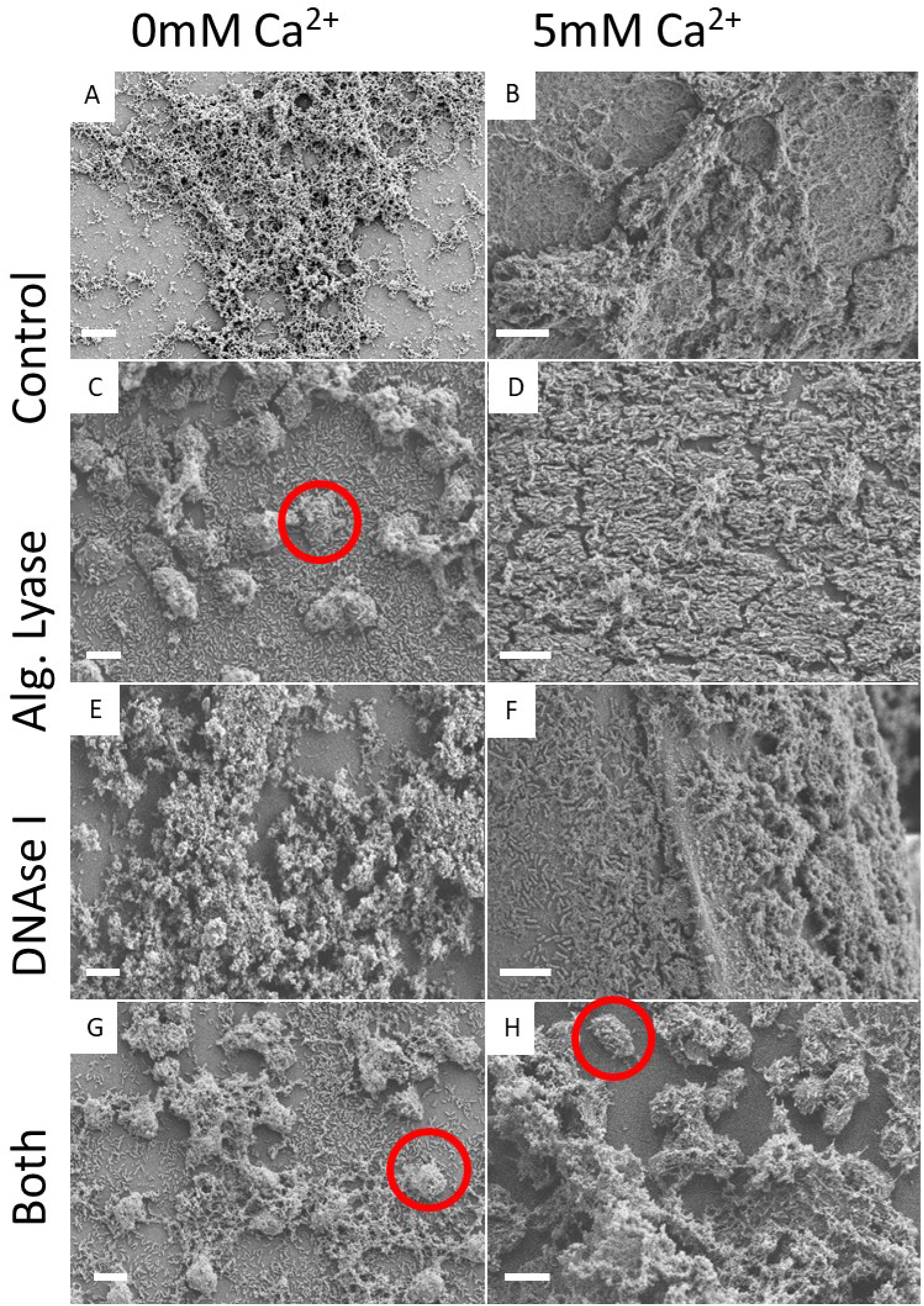
Large-scale structure of biofilms grown with and without 5mM Ca^2+^ after targeted enzyme treatments. All biofilms have been incubated with neutrophils for 1 hour prior to fixation. A) An untreated biofilm without Ca^2+^ contains large-scale alginate structures. B) With Ca^2+^, biofilms exhibit more substrate surface coverage. C) When treated with alginate lyase, the alginate structures in biofilms without Ca^2+^ have been compromised. Neutrophils (example circled in red) and planktonic bacterial cells are present on the substrate D) Biofilms with Ca^2+^ treated with alginate lyase exhibit a large-scale structure of closely-packed, parallel-aligned bacterial cells along the substrate. A false-colored image of this panel has been reproduced as Supplementary Fig. 4 E) No substantial changes are observed for biofilms without Ca^2+^ treated with DNAse I. F) When treated with DNAse I, biofilms with Ca^2+^ no longer present the aligned structure seen in Panel D. G) Biofilms without Ca^2+^ treated with both enzymes and H) biofilms with Ca^2+^ treated with both enzymes each exhibit large-scale structural compromise and the prominent presence of neutrophils (example circled in red). Scale bars represent 10 µm.

Compared to biofilms grown without added Ca^2+^ (Figure 4A), those grown with 5mM Ca^2+^ (Figure 4B) exhibit more surface coverage of the substrate and a more contiguous structure. This is consistent with our observations of a thicker biofilm with a dense, contiguous layer of DNA shown in Figure 2B,D.

For biofilms without added Ca^2+^ and treated with alginate lyase (Figure 4C), we observe large-scale structural compromise indicated by substantially smaller, more discrete pieces of alginate and an increase of exposed single cells of bacteria on the substrate. When biofilms are grown with 5mM Ca^2+^ and then treated with alginate lyase we observe a layer of closely-packed, aligned bacterial cells along the substrate (Figure 4D). As this structure does not appear to be compromised by alginate lyase, we hypothesize that eDNA is the material binding bacterial cells together. This inference is supported by the layer of DNA shown in Figure 2D.

When biofilms are grown without Ca^2+^ and treated with DNAse I, the biofilm structure does not appear to be substantially altered from the untreated case (Figure 4E). For biofilms grown with 5mM Ca^2+^ and treated with DNAse I, we observe an increase of non-aligned single bacterial cells near the surface of the substrate, indicating that the structure observed in Figure 4D has been compromised by treatment with DNAse I. From this, we infer that the shield-like structure in Figures 3D and 4D is mediated by eDNA.

When biofilms are grown without Ca^2+^ and treated with both enzymes (Figure 4G), the structural changes appear similar to those caused by treatment with alginate lyase alone (Figure 4C). In contrast, when biofilms are grown with 5mM Ca^2+^ and treated with a combination of both enzymes, (Figure 4H), the alginate structures are broken up and the sheet-like architecture of aligned cells (Figure 4D) is gone entirely.

### The interactions of neutrophils with biofilms are differentially impacted by enzyme treatments

The ∼10 μm round objects seen in Figure 4 C, G, and H are neutrophils. Notably, though all samples have been incubated with neutrophils, their prominent presence is only observed for biofilms grown without Ca^2+^ and treated with alginate lyase (Figure 4C) and for biofilms grown either without or with Ca^2+^ and then treated with a combination of both enzymes (Figure 4G,H). Since all samples had the same period of incubation with the same concentration of neutrophils, we interpret these findings as indicating that neutrophils are much better retained in these three types of samples throughout the subsequent washing and fixation process. We further attribute the retention of neutrophils to their successful interactions with the biofilm, which corresponds with increases in phagocytic success (Figure 1A,B). The biofilm is attached to the substrate and the neutrophils are not; those neutrophils unable to establish a successful involvement with biofilm material may have been washed away during sample preparation.

The absence of neutrophils in SEM images biofilms grown with 5mM Ca^2+^ and treated with alginate lyase (Figure 4D) suggests that the layer of tightly-packed bacteria, which is many times larger than the diameter of a neutrophil, may help protect against phagocytosis. Treatment with alginate lyase reveals this underlying structure but does not result in increased phagocytic success (Figure 1B). Indeed, for biofilms grown with 5mM Ca^2+^, only the combination of alginate lyase and DNAse I results in compromise of both structure types and substantial neutrophil presence in SEM images (Figure 4H) as well as a substantial increase in engulfment success (Figure 1B).

Figures 2D, 3B, and 4B show that biofilms grown with Ca^2+^ have large-scale structures that are compositionally and microstructurally distinct from the rest of the biofilm and that are not found in biofilms grown without Ca^2+^. These structures are broad, plate-like aggregates of tightly-packed bacteria connected by a web of eDNA. Within these plates, bacteria are largely aligned parallel to one another. Alignment of bacteria into aggregates has previously been associated with depletion attraction [49, 50]. However, here the size and 2-D, planar shape of the aggregates, their proximity to the substrate, the observation that eDNA is binding cells together, and this structure’s dependence on supplementary Ca^2+^, are all indicative that these planar “shields” arise from something other than depletion attraction.

False-colored versions of Figure 3D and Figure 4D are included as Supplementary Figures S3 and S4. False coloring is done to distinguish the regions of alginate from the tightly-packed bacterial shield, and to make more visible the eDNA that binds bacteria together to make the shield.

Previous work by others has demonstrated that eDNA facilitates surface colonization and motility [48, 51, 52]. Divalent calcium cations specifically have been shown to interact with DNA at larger length scales and strengths than singly-charged sodium cations [48], which are also present in bacterial media and *in vivo*. We speculate that this strong cationic influence of Ca^2+^ augments the eDNA’s polymer bridging capacity to produce the distinct microdomains we find in biofilms grown in the presence of Ca^2+^. We also show that this “shield” microstructure plays a protective role by inhibiting phagocytosis if the larger alginate structure becomes compromised.

Free Ca^2+^ ions are present in CF lungs in the range of 1-5mM. Thus, the new understanding developed in the present work will help to advance treatments of chronic *P. aeruginosa* infections in CF airways, which are largely caused by alginate-producing strains. We propose that the ability to eradicate these biofilm infections via mechanical and microstructural compromise must necessarily rely on knowledge of the biofilm’s spatial organization and regional architectures. Targeting a single specific EPS component, such as alginate, without full characterization of the biofilm, has an insignificant effect on enhancing clearance by neutrophils, and may in fact exacerbate the immune response to infection [14, 53]. In contrast, designing multi-component treatment plans that maximize structural disruption and promote success of the body’s natural responses may expedite biofilm eradication and hinder the evolutionary development of antibiotic resistance by reducing amount of antibiotic needed in treatment.

For the work presented here in vitro, an alginate-dominant strain of *P. aeruginosa* was used to mimic the mucoid phenotypes found in CF lungs [5, 44, 54, 55]. In addition to alginate and eDNA, *P. aeruginosa* produces the polysaccharides Pel and Psl. Pel, which is cationic, has been shown to cross-link with eDNA in in the biofilm matrix [56] and increase eDNA’s resistance to digestion by DNAse I [28, 49]. As such, biofilms containing significant amounts of Pel may further hinder the compromise of protective regional structures in the EPS matrix. Future studies are needed to investigate the extent of the phenomena observed in this work in the presence of other self-produced and host-derived EPS components.

Our results demonstrate that Ca^2+^ facilitates the formation of compositionally and structurally distinct regions. Others have previously shown that Ca^2+^ cross-links alginate to form a gel-like biofilm in which bacteria are diffusely distributed [44] and that Ca^2+^ mediates eDNA-based aggregation of bacterial cells [51]. We have shown the occurrence of both phenomena simultaneously within Ca^2+^- gelled mucoid biofilms, leading to the formation of distinct organizational layers. Further, the presence of regional structures separately dominated by either alginate or eDNA serves as a protective mechanism against phagocytosis by neutrophils. When either alginate or eDNA is targeted individually with polymer-degrading enzymes, the remaining matrix microstructure is sufficient to protect the biofilm bacteria from phagocytosis by neutrophils.

The implications of the findings reported here are significant in the study of prevention and control of biofilm disease. Strategies to combat biofilm growth include anti-biofilm coatings to repel bacteria with bactericidal, topographical, chemical, or electrostatic interactions [57-66]. Indeed, one study revealed that modifying surfaces with immobilized DNA leads to reduced bacterial attachment, potentially due to electrostatic repulsion between the negative charges of DNA and bacterial cell surfaces [67]. The clinical implementation of DNA-coated surfaces may necessarily rely on an understanding of ionic interactions at the biofilm-substrate interface. The work presented here highlights the effects of metal ions incorporated from the host environment on internal biofilm organization, an added layer of complexity beyond the biofilm-substrate interface.

## Conclusion

Here we have investigated the effects of targeted enzyme treatment on promoting phagocytosis of *P. aeruginosa* from mucoid biofilms. For biofilms grown *in vitro* without Ca^2+^, we correlate phagocytic success with decreases in the relative elasticity of biofilms and compromise of structural organization. Our results show that biofilms grown under these conditions are primarily protected by alginate in their EPS matrix. Treatment with alginate lyase results in both a shift toward lower relative elasticity and an increase in phagocytic success, while DNAse I has no significant effect on either quantity (Figure 1 A,C). However, when biofilms are grown with 5mM Ca^2+^, treatment with either alginate lyase or DNAse I decreases relative elasticity, but only treatment with the enzymes in combination increases phagocytic success (Figure 1 B,D).

Confocal laser-scanning microscopy reveals the presence of small, discrete regions of eDNA in biofilms grown without added calcium. In biofilms grown with 5mM Ca^2+^ we observe a dense, contiguous, flat structure of eDNA (Figure 2D).

Corresponding with this plane of dense eDNA, scanning electron microscopy provides detailed images of a sheet-like layer of closely aligned bacterial cells bound by a web of eDNA (Figure 3 B). This structure survives treatment with alginate lyase and is only compromised by DNAse I. Substantial structural compromise of the biofilm requires a combination enzyme treatment.

### Supporting Information

Ensemble averaged MSD-lag time plots from microrheology expreiments; green fluorescence of biomass from biofilms, imaged with confocal laser scanning microscopy; false-colored reproduction of Figure 3D; false-colored reproduction of Figure 4D; representative images of neutrophils assessed for phagocytic success for each growth condition and treatment type; table of CFU/mL for each biofilm type in biological triplicate.

## Acknowledgements

SEM preparation and imaging was performed at the Center for Biomedical Research

Support Microscopy and Imaging Facility at UT Austin (RRID:SCR_021756).

## Supplementary figures

**Figure S1.**
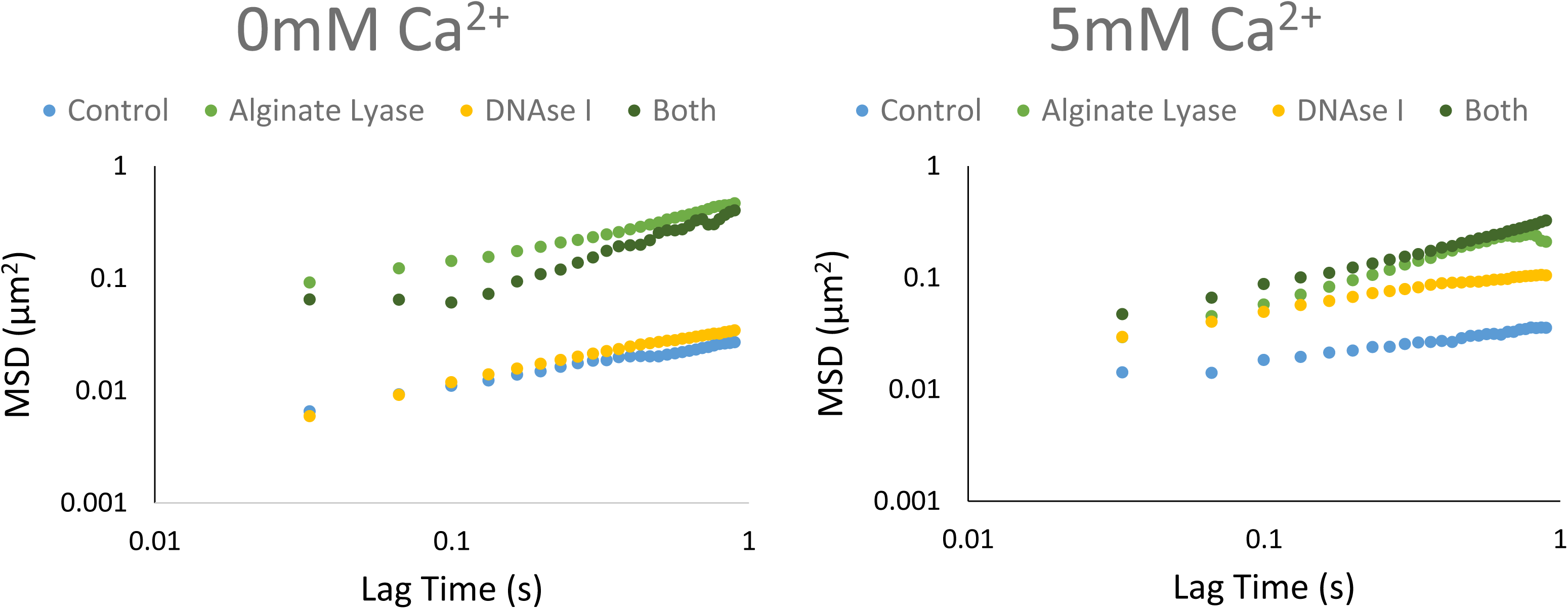
Ensemble averaged MSD-lag time plots from experiments with biofilms with and without added Ca2+ and each enzyme treatment type

**Figure S2.**
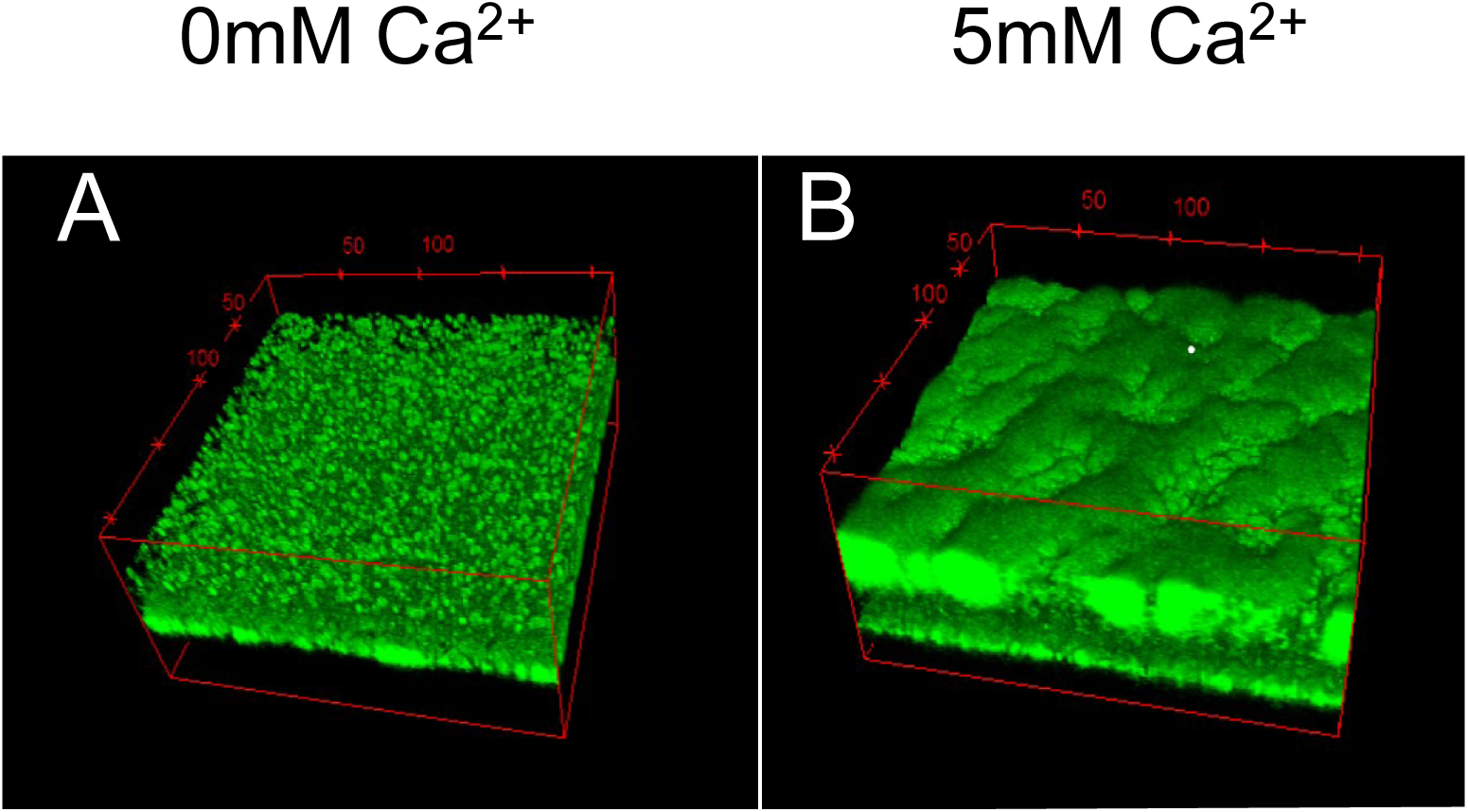
Green fluorescence of biomass from biofilms, imaged with confocal laser scanning microscopy. Coordinate boxes have dimensions of 212×212×100µm.

**Figure S3.**
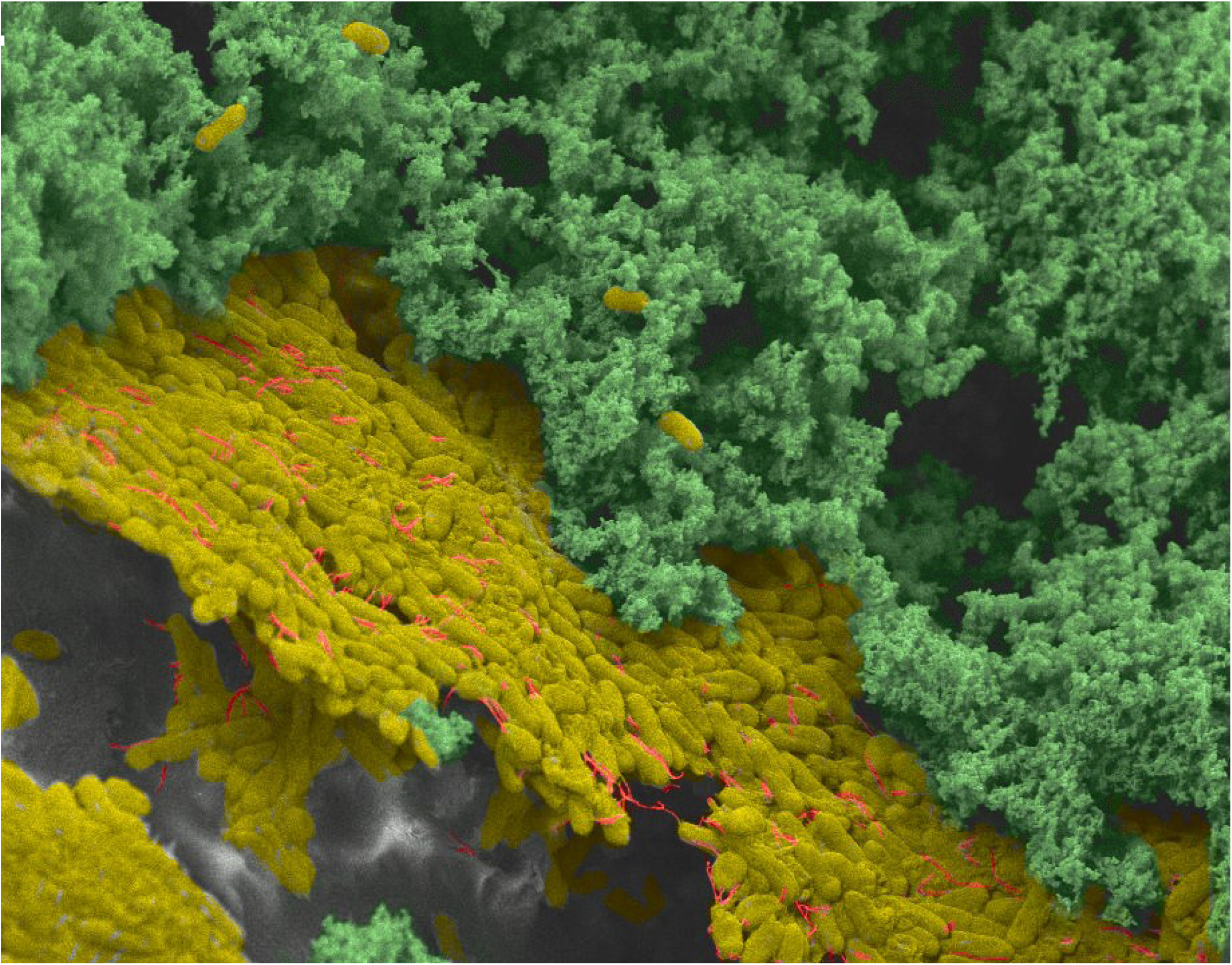
False-color reproduction of Figure 3B. Bacteria are yellow in yellow, alginate in green, and eDNA in red. The scale bar represents 2 μm

**Figure S4.**
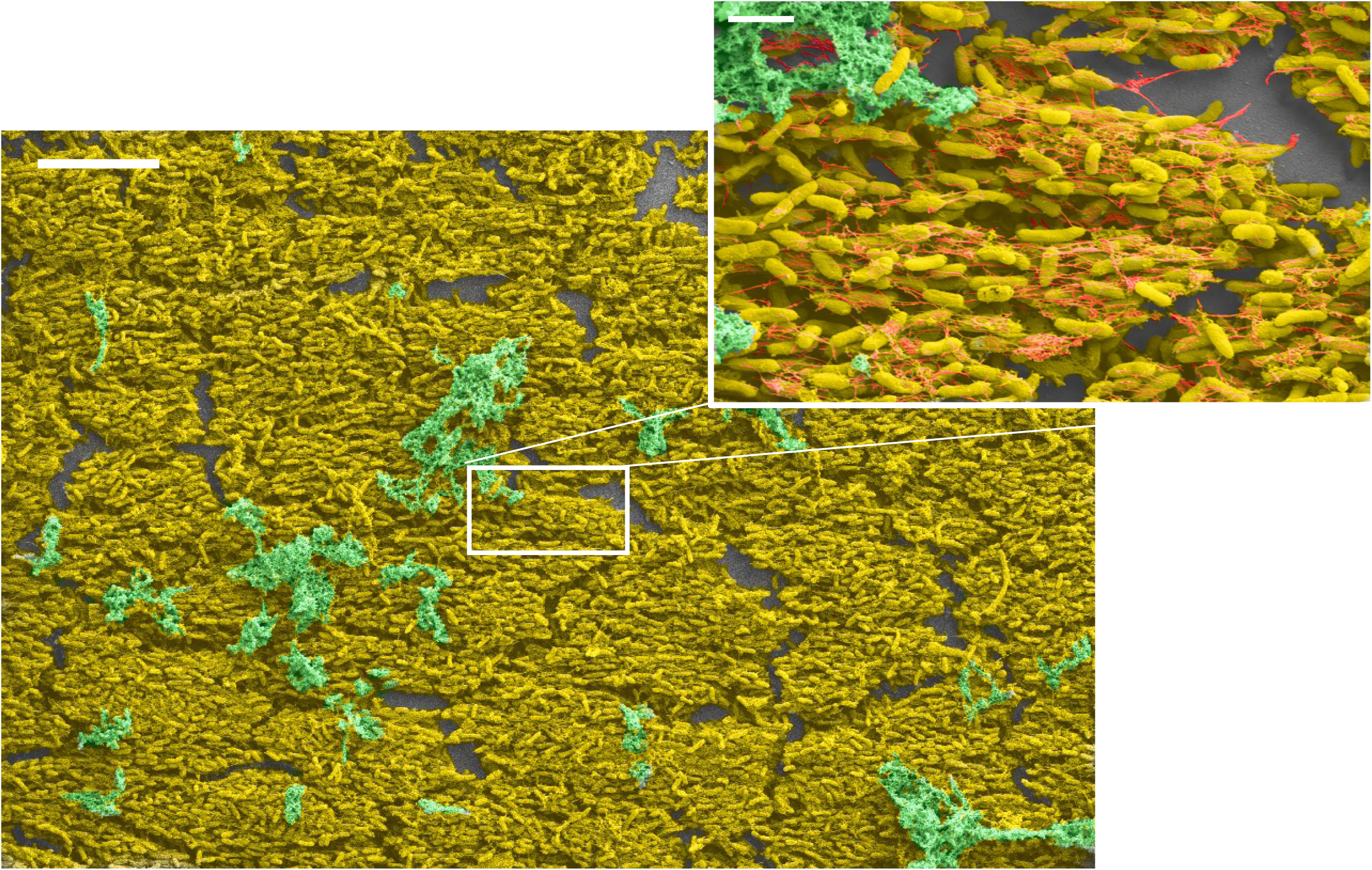
False-colored reproduction of the large-scale structure of parallel bacterial cells shown in Figure 4D. Bacteria are represented in yellow, alginate in green, and eDNA in red. The inset shows a higher magnification of the selected region. The scale bars represent 10 μm for the larger image and 2 μm for the inset.

**Figure S5.**
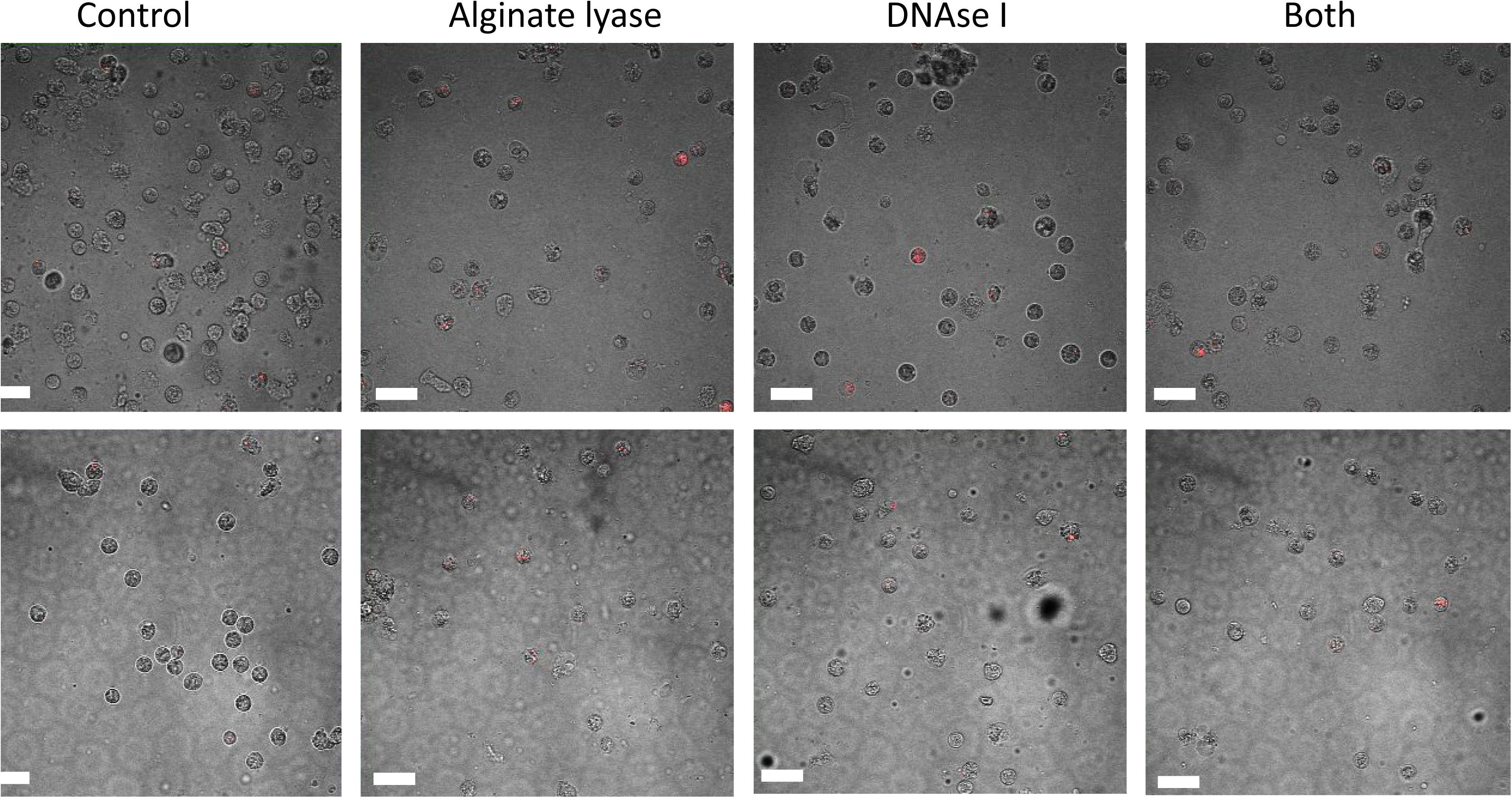
Representative images of neutrophils assessed for phagocytic success for each biofilm and treatment combination. Bacteria have been stained with pHrodo (ThermoFisher Scientific), which fluoresces red only within the acidic phagosome. Scale bars represent 20 μm

## References

1. Blanchard, A.C. and V.J. Waters, Opportunistic Pathogens in Cystic Fibrosis: Epidemiology and Pathogenesis of Lung Infection. J Pediatric Infect Dis Soc, 2022. 11(Supplement_2): p. S3–S12.

2. Costerton, J.W., P.S. Stewart, and E.P. Greenberg, Bacterial biofilms: a common cause of persistent infections. Science, 1999. 284(5418): p. 1318–22.

3. Martin, I., V. Waters, and H. Grasemann, Approaches to Targeting Bacterial Biofilms in Cystic Fibrosis Airways. Int J Mol Sci, 2021. 22(4).

4. Planet, P.J., Adaptation and Evolution of Pathogens in the Cystic Fibrosis Lung. J Pediatric Infect Dis Soc, 2022. 11(Supplement_2): p. S23–S31.

5. Høiby, N., O. Ciofu, and T. Bjarnsholt, Pseudomonas aeruginosa biofilms in cystic fibrosis. Future Microbiol, 2010. 5(11): p. 1663–74.

6. Birkenhauer, E., S. Neethirajan, and J.S. Weese, Collagen and hyaluronan at wound sites influence early polymicrobial biofilm adhesive events. BMC Microbiol, 2014. 14: p. 191.

7. Papayannopoulos, V., Neutrophils Facing Biofilms: The Battle of the Barriers. Cell Host Microbe, 2019. 25(4): p. 477–479.

8. Hills, O.J., et al., Cation complexation by mucoid Pseudomonas aeruginosa extracellular polysaccharide. PLoS One, 2021. 16(9): p. e0257026.

9. Kovach, K., et al., Evolutionary adaptations of biofilms infecting cystic fibrosis lungs promote mechanical toughness by adjusting polysaccharide production. NPJ Biofilms Microbiomes, 2017. 3: p. 1.

10. Murgia, X., et al., Micro-rheological properties of lung homogenates correlate with infection severity in a mouse model of Pseudomonas aeruginosa lung infection. Scientific Reports, 2020. 10(1): p. 16502.

11. Segal, A.W., How neutrophils kill microbes. Annu Rev Immunol, 2005. 23: p. 197–223.

12. Brinkmann, V., et al., Neutrophil extracellular traps kill bacteria. Science, 2004. 303(5663): p. 1532–5.

13. Pérez-Figueroa, E., et al., Neutrophils: Many Ways to Die. Front Immunol, 2021. 12: p. 631821.

14. Rada, B., Interactions between Neutrophils and Pseudomonas aeruginosa in Cystic Fibrosis. Pathogens, 2017. 6(1).

15. Mulcahy, H., L. Charron-Mazenod, and S. Lewenza, Extracellular DNA chelates cations and induces antibiotic resistance in Pseudomonas aeruginosa biofilms. PLoS Pathog, 2008. 4(11): p. e1000213.

16. Olivares, E., et al., Clinical Impact of Antibiotics for the Treatment of. Front Microbiol, 2019. 10: p. 2894.

17. Tseng, B.S., et al., The extracellular matrix protects Pseudomonas aeruginosa biofilms by limiting the penetration of tobramycin. Environ Microbiol, 2013. 15(10): p. 2865–78.

18. Mah, T.F. and G.A. O’Toole, Mechanisms of biofilm resistance to antimicrobial agents. Trends Microbiol, 2001. 9(1): p. 34–9.

19. Rahman, M.U., et al., Effect of collagen and EPS components on the viscoelasticity of. Soft Matter, 2021. 17(25): p. 6225–6237.

20. Eichelberger, K.R. and W.E. Goldman, Manipulating neutrophil degranulation as a bacterial virulence strategy. PLoS Pathog, 2020. 16(12): p. e1009054.

21. Gloag, E.S., et al., Biofilm mechanics: Implications in infection and survival. Biofilm, 2020. 2: p. 100017.

22. Gordon, V.D.D.-F., Megan Kovach, Kristin Rodesney, Christopher A, Biofilms and Mechanics: A Review of Experimental Techniques and Findings. Journal of Physics D: Applied Physics, 2017. 50(22).

23. Stewart, P.S., Biophysics of biofilm infection. Pathog Dis, 2014. 70(3): p. 212–8.

24. Wells, M., et al., Perspective: The viscoelastic properties of biofilm infections and mechanical interactions with phagocytic immune cells. Front Cell Infect Microbiol, 2023. 13: p. 1102199.

25. Rahman, M.U., et al., Microrheology of Pseudomonas aeruginosa biofilms grown in wound beds. NPJ Biofilms Microbiomes, 2022. 8(1): p. 49.

26. Davis-Fields, M., et al., Assaying How Phagocytic Success Depends on the Elasticity of a Large Target Structure. Biophys J, 2019. 117(8): p. 1496–1507.

27. Bakhtiari, L.A., M.J. Wells, and V.D. Gordon, High-throughput assays show the timescale for phagocytic success depends on the target toughness. Biophys Rev (Melville), 2021. 2(3): p. 031402.

28. Kovach, K.N., et al., Specific Disruption of Established Pseudomonas aeruginosa Biofilms Using Polymer-Attacking Enzymes. Langmuir, 2020. 36(6): p. 1585–1595.

29. Fleeman, R.M., M. Mikesh, and B.W. Davies, Investigating Klebsiella pneumoniae biofilm preservation for scanning electron microscopy. Access Microbiol, 2023. 5(2).

30. Schindelin, J., et al., Fiji: an open-source platform for biological-image analysis. Nat Methods, 2012. 9(7): p. 676–82.

31. Mason, T.G., et al., Particle Tracking Microrheology of Complex Fluids. Physical Review Letters, 1997. 79(17): p. 3282–3285.

32. Mason, T.G. and D.A. Weitz, Optical Measurements of Frequency-Dependent Linear Viscoelastic Moduli of Complex Fluids. Physical Review Letters, 1995. 74(7): p. 1250–1253.

33. Savin, T. and P.S. Doyle, Statistical and sampling issues when using multiple particle tracking. Physical Review E, 2007. 76(2): p. 021501.

34. Tinevez, J.Y., et al., TrackMate: An open and extensible platform for single-particle tracking. Methods, 2017. 115: p. 80–90.

35. Tarantino, N., et al., TNF and IL-1 exhibit distinct ubiquitin requirements for inducing NEMO-IKK supramolecular structures. J Cell Biol, 2014. 204(2): p. 231–45.

36. Zhu, B. and H. Yin, Alginate lyase: Review of major sources and classification, properties, structure-function analysis and applications. Bioengineered, 2015. 6(3): p. 125–31.

37. Hatch, R.A. and N.L. Schiller, Alginate lyase promotes diffusion of aminoglycosides through the extracellular polysaccharide of mucoid Pseudomonas aeruginosa. Antimicrob Agents Chemother, 1998. 42(4): p. 974–7.

38. Islan, G.A., V.E. Bosio, and G.R. Castro, Alginate lyase and ciprofloxacin co-immobilization on biopolymeric microspheres for cystic fibrosis treatment. Macromol Biosci, 2013. 13(9): p. 1238–48.

39. Patel, K.K., et al., Alginate lyase immobilized chitosan nanoparticles of ciprofloxacin for the improved antimicrobial activity against the biofilm associated mucoid P. aeruginosa infection in cystic fibrosis. Int J Pharm, 2019. 563: p. 30–42.

40. Converter between molecular weight (kDa) and hydrodynamic radius (nm). 2023; Available from: https://www.fluidic.com/toolkit/hydrodynamic-radius-converter/.

41. Teirlinck, E., et al., Functionalized Nanomaterials for the Management of Microbial Infection. 2017, Elsevier. p. 49–76.

42. Chew, S.C., et al., Dynamic remodeling of microbial biofilms by functionally distinct exopolysaccharides. mBio, 2014. 5(4): p. e01536–14.

43. Cohen-Cymberknoh, M., et al., Calcium carbonate mineralization is essential for biofilm formation and lung colonization. iScience, 2022. 25(5): p. 104234.

44. Jacobs, H.M., et al., Mucoid Pseudomonas aeruginosa Can Produce Calcium-Gelled Biofilms Independent of the Matrix Components Psl and CdrA. J Bacteriol, 2022. 204(5): p. e0056821.

45. Das, T., et al., Influence of calcium in extracellular DNA mediated bacterial aggregation and biofilm formation. PLoS One, 2014. 9(3): p. e91935.

46. Sarkisova, S., et al., Calcium-induced virulence factors associated with the extracellular matrix of mucoid Pseudomonas aeruginosa biofilms. J Bacteriol, 2005. 187(13): p. 4327–37.

47. Kerchove, A.J. and M. Elimelech, Calcium and magnesium cations enhance the adhesion of motile and nonmotile pseudomonas aeruginosa on alginate films. Langmuir, 2008. 24(7): p. 3392–9.

48. Morales-García, A.L., et al., The Role of Extracellular DNA in Microbial Attachment to Oxidized Silicon Surfaces in the Presence of Ca. Langmuir, 2021. 37(32): p. 9838–50.

49. Jennings, L.K., et al., Pseudomonas aeruginosa aggregates in cystic fibrosis sputum produce exopolysaccharides that likely impede current therapies. Cell Rep, 2021. 34(8): p. 108782.

50. Secor, P.R., et al., Entropically driven aggregation of bacteria by host polymers promotes antibiotic tolerance in. Proc Natl Acad Sci U S A, 2018. 115(42): p. 10780–10785.

51. Gloag, E.S., et al., Self-organization of bacterial biofilms is facilitated by extracellular DNA. Proc Natl Acad Sci U S A, 2013. 110(28): p. 11541–6.

52. Panlilio, H. and C.V. Rice, The role of extracellular DNA in the formation, architecture, stability, and treatment of bacterial biofilms. Biotechnol Bioeng, 2021. 118(6): p. 2129–2141.

53. Walker, T.S., et al., Enhanced Pseudomonas aeruginosa biofilm development mediated by human neutrophils. Infect Immun, 2005. 73(6): p. 3693–701.

54. Pedersen, S.S., et al., Role of alginate in infection with mucoid Pseudomonas aeruginosa in cystic fibrosis. Thorax, 1992. 47(1): p. 6–13.

55. Hentzer, M., et al., Alginate overproduction affects Pseudomonas aeruginosa biofilm structure and function. J Bacteriol, 2001. 183(18): p. 5395–401.

56. Jennings, L.K., et al., Pel is a cationic exopolysaccharide that cross-links extracellular DNA in the Pseudomonas aeruginosa biofilm matrix. Proc Natl Acad Sci U S A, 2015. 112(36): p. 11353–8.

57. Li, P., et al., Bacterial Biofilm Formation on Biomaterials and Approaches to Its Treatment and Prevention. Int J Mol Sci, 2023. 24(14).

58. Samuel, M.S., et al., A Flexible Anti-Biofilm Hygiene Coating for Water Devices. ACS Appl Bio Mater, 2022. 5(8): p. 3991–3998.

59. Ishihama, H., et al., An antibacterial coated polymer prevents biofilm formation and implant-associated infection. Sci Rep, 2021. 11(1): p. 3602.

60. Jahanmard, F., et al., Bactericidal coating to prevent early and delayed implant-related infections. J Control Release, 2020. 326: p. 38–52.

61. Ahmadabadi, H.Y., K. Yu, and J.N. Kizhakkedathu, Surface modification approaches for prevention of implant associated infections. Colloids Surf B Biointerfaces, 2020. 193: p. 111116.

62. Piras, A.M., et al., *Antibacterial, Antibiofilm,* and Antiadhesive Properties of Different Quaternized Chitosan Derivatives. Int J Mol Sci, 2019. 20(24).

63. Hoque, J., et al., Charge-Switchable Polymeric Coating Kills Bacteria and Prevents Biofilm Formation in Vivo. ACS Appl Mater Interfaces, 2019. 11(42): p. 39150–39162.

64. Acosta, S., et al., Antibiofilm coatings based on protein-engineered polymers and antimicrobial peptides for preventing implant-associated infections. Biomater Sci, 2020. 8(10): p. 2866–2877.

65. Yi, S.Y., et al., Substrate-independent adsorption of nanoparticles as anti-biofilm coatings. Biomater Sci, 2022. 10(2): p. 410–422.

66. Yadav, N., et al., Graphene Oxide-Coated Surface: Inhibition of Bacterial Biofilm Formation due to Specific Surface-Interface Interactions. ACS Omega, 2017. 2(7): p. 3070–3082.

67. Pingle, H., et al., Minimal attachment of. Biointerphases, 2018. 13(6): p. 06E405.

