## supplementary material for "Physiological concentrations of calcium interact with alginate and extracellular DNA in the matrices of *Pseudomonas aeruginosa* biofilms to impede phagocytosis by neutrophils"

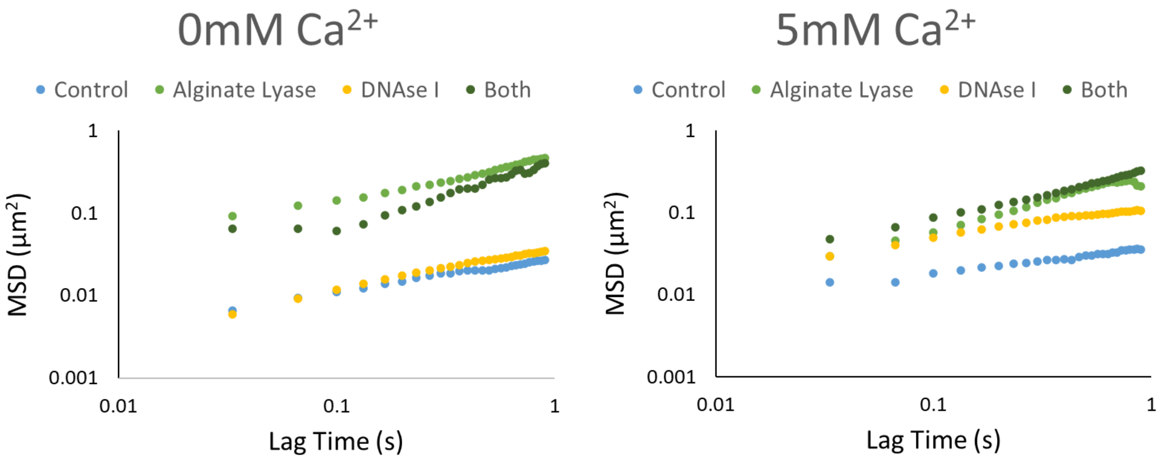


Figure S1. Ensemble averaged MSD-lag time plots from experiments with biofilms with and without added $Ca^{2+}$ and each enzyme treatment type

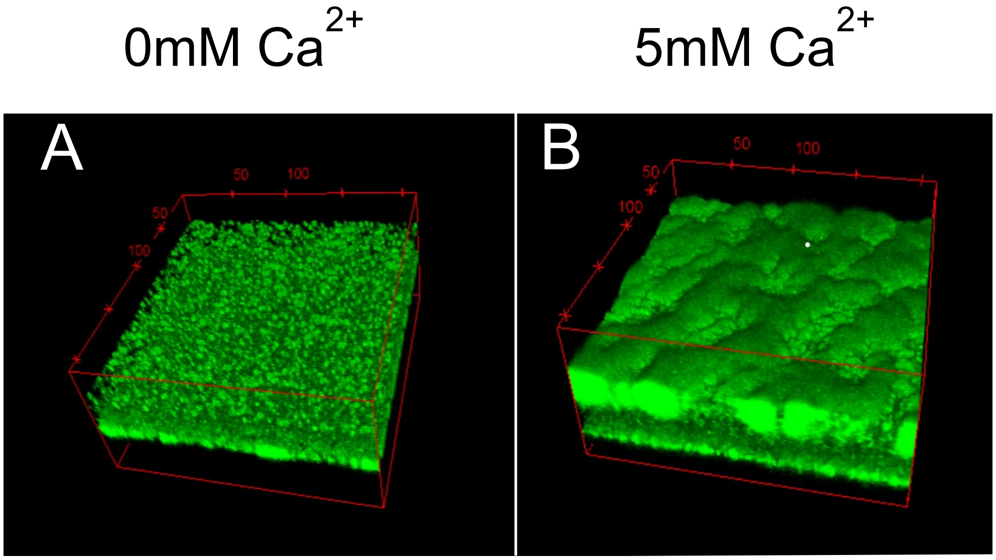


Figure S2. Green fluorescence of biomass from biofilms, imaged with confocal laser scanning microscopy. Coordinate boxes have dimensions of 212x212x100µm.

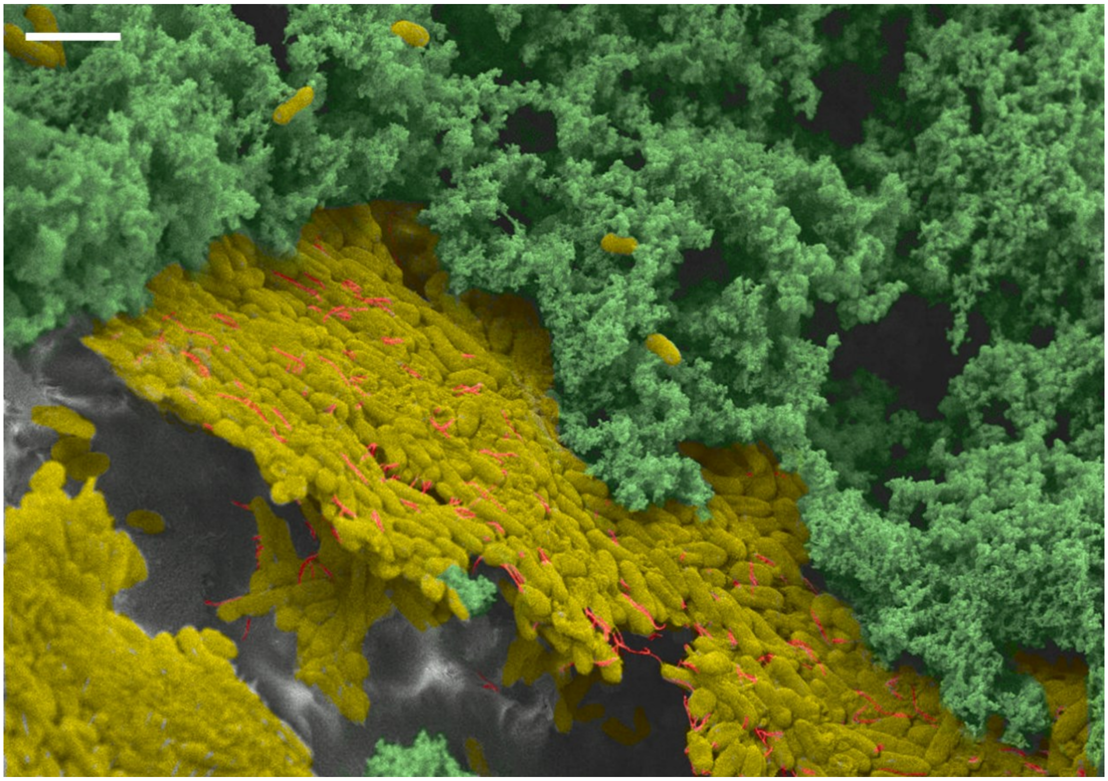

Figure S3. False-color reproduction of Figure 3D. Bacteria are yellow in yellow, alginate in green, and eDNA in red. The scale bar represents 2 µm

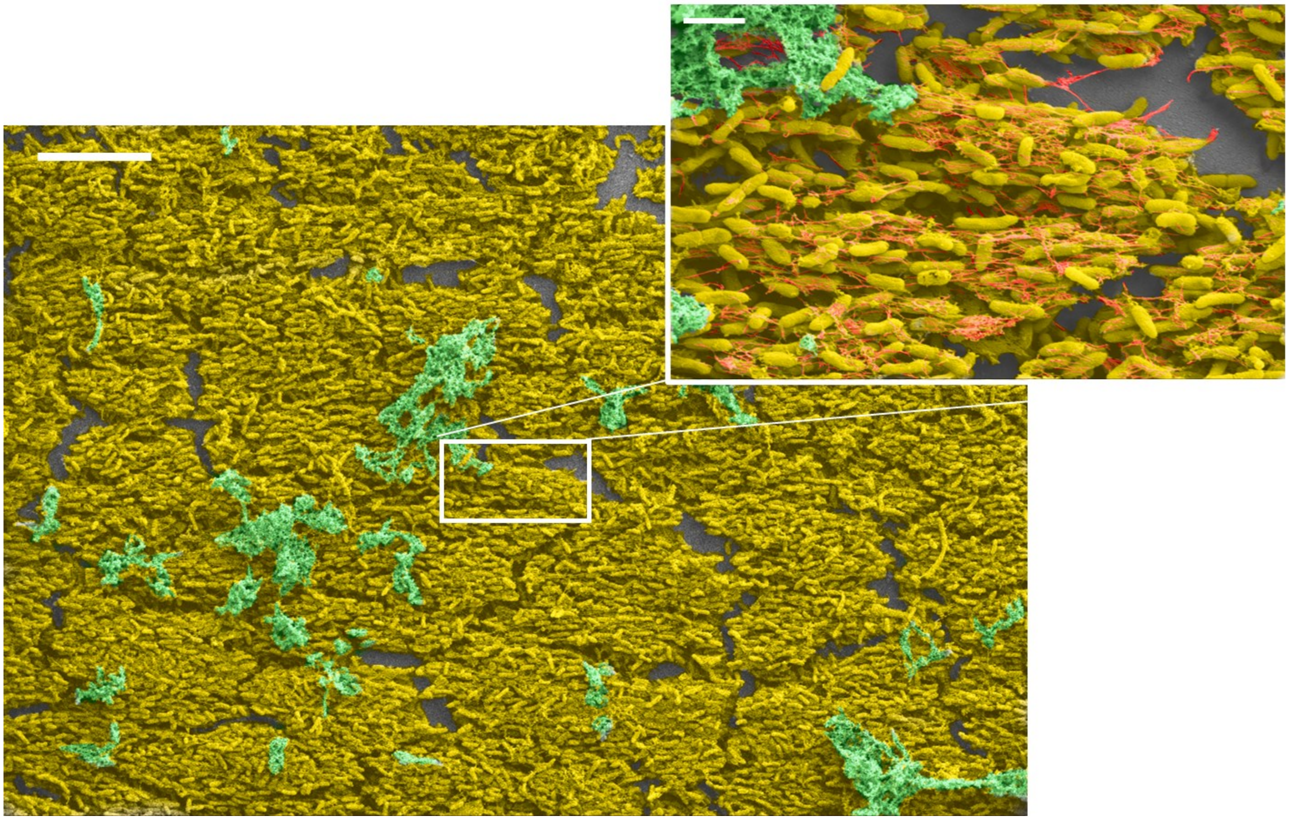

Figure S4. False-colored reproduction of the large-scale structure of parallel bacterial cells shown in Figure 4D. Bacteria are represented in yellow, alginate in green, and eDNA in red. The inset shows a higher magnification of the selected region. The scale bars represent 10 µm for the larger image and 2 µm for the inset.

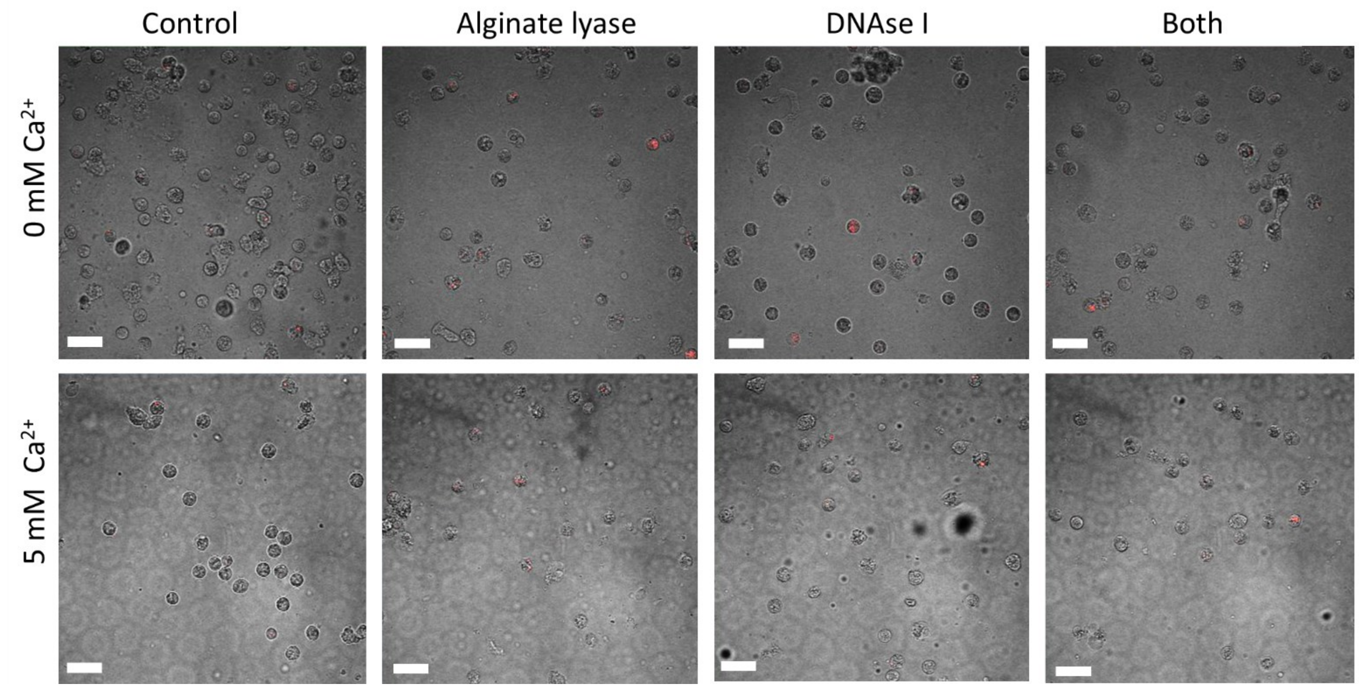


Figure S5. Representative images of neutrophils assessed for phagocytic success for each biofilm and treatment combination. Bacteria have been stained with pHrodo (ThermoFisher Scientific), which fluoresces red only within the acidic phagosome. Scale bars represent 20 µm.

| | 0mM $Ca^{2+}$ | 5mM $Ca^{2+}$ | Statistics | |
|---|---|---|---|---|
| Replicate | CFU/mL | CFU/mL | ANOVA p-value | Student t-test p-value |
| 1 | $1.92x10^4$ | $1.95x10^5$ | 0.204 | 0.204 |
| 2 | $7.20x10^4$ | $2.40x10^4$ | | |
| 3 | $2.14x10^4$ | $4.15x10^5$ | | |

Table S1. Colony-forming units from biofilms grown without and with 5mM $Ca^{2+}$. ANOVA and Student t-tests yielded no significance between the groups. Experiments were performed in biological triplicate.
